# Butterfly body size shrinkage: the impact of ecological traits across varied environments

**DOI:** 10.1101/2025.02.06.636804

**Authors:** Konstantina Zografou, Eva Knop, Brent J Sewall, Oliver Schweiger, Consuelo M De Moraes, Melissa Whitaker, Vassiliki Kati

## Abstract

Because of its close ties to numerous ecological and life history characteristics, body size is regarded as one of an organism’s most important characteristics and is frequently considered a significant indicator of fitness. According to recent studies, ectotherms in particular, may see a reduction in body size with rising temperatures. How life history and ecological traits influence, however, shifts in butterfly body size in response to environmental changes, particularly focusing on the effects of temperature and land use is poorly studied. Using Generalized Additive Models (GAM), we analyzed forewing length data alongside various life history (phenology, overwintering developmental stage, voltinism, diet breadth, gender) and ecological traits (mobility, thermal tolerance) as well as environmental parameters for a period that lasts 110 years. Smaller body sizes were linked to early-season emergence and increasing forest cover, while larger sizes were linked to longer flight durations and later seasonal appearance. Males exhibited a pronounced decline in body size while females showed an opposite trend, suggesting sex-specific vulnerabilities to climate change. This research highlights the complex interplay between climate change, habitat fragmentation, and butterfly morphology, emphasizing the need for further investigation into sexual size dimorphism as anthropogenic influences continue to reshape butterfly populations.

## Introduction

Climate change is a pressing global challenge that significantly impacts biodiversity (Garcia et al. 2014)and in particular ectothermic organisms, like butterflies. Butterflies are highly sensitive to anthropogenic changes and due to their rapid reaction time, they often serve as effective indicators of ecosystem health (Parmesan 2003, Hill et al. 2021). These organisms exhibit a gradient of life history traits, including but not limited to phenology, overwintering stages, voltinism, diet, and gender, which mold their responses to changing environmental conditions (Fischer and Fiedler 2002, Altermatt 2010, Karl et al. 2011, Zografou et al. 2021). Other than the life history traits, ecological traits such as geographical distribution range and thermal tolerance, and physiological traits such as body size may determine species adaptation to climate change. The time of occurrence of life cycle events (phenology) is one of the most sensitive biological traits to climate change. The difference in ecological traits can clearly indicate different phenological responses; for example, multivoltine species (with more than two generations per year) have flying periods extended and reproduce more generations compared with the single-generating species (univoltine), which allow them to adapt quickly to warmer temperatures (Zhang et al., 2021). Furthermore, an important trait that influences organisms’ resilience to changing climate is dietary preferences. There are polyphagous species that can utilize a wider range of host plants and thus capable of overcoming limitations in nutrition because of phenological mismatches between herbivores and their hosts (Singer and Parmesan 2010) and monophagous, with strictly specialized diets (Scheffers et al. 2016).

Mobility and thermal tolerance are ecological traits that regulate butterflies’ adaptation to changing temperatures. Highly mobile species will be able to shift their geographical range to suitable habitats more successfully than their less mobile counterparts (Hoffmann and Sgrò 2011). For instance, increased mobility has been proved to help organisms to distribute within urbanized, highly fragmented areas (Merckx et al. 2018). In addition, butterflies having high thermal tolerance can sustain extreme heat events (Kingsolver et al. 2013) which is vital for survival as temperatures rise. Another morphological trait that is physiologically relevant includes forewing length (a proxy for the body size), which is related to thermoregulation and dispersal ability. In theory, larger wings may benefit flying performance; however, wing sizes are also influenced by environmental conditions. For instance, species temperature rule states that when species are raised in warmer temperatures tend to decrease adult body sizes (Angilletta and Dunham 2003). Changes in land use have a substantial impact on these ecological and life history characteristics. Local temperatures and resource availability can be influenced by the unique microhabitats created by various land use types including pastures, urban areas, woodlands, and agricultural crops. For instance, urban regions can cause habitat fragmentation and higher levels of heat stress in populations of butterflies.

Merckx et al. (2018) found that urban-heat-island effect decreased the size of numerous taxa except for two ectotherm groups. However, substantial habitat fragmentation and the positive covariation between size and dispersal led to larger body sizes for butterflies and Orthoptera.

Linkages among life history traits and land use cover in the context of climate change are known to have an increasingly important role (Altermatt 2012, Oliver and Morecroft 2014). However, how these linkages influence body size of butterflies remain poorly studied. As butterflies adapt their phenology and morphology in response to changing climate and land use cover, understanding these interactions will improve our ability to forecast future population dynamics.

In this study, we build upon previous findings that indicate a decrease in forewing length among butterfly populations over the past century (*unpublished data*). Specifically, we ask two primary research questions: 1) How do life history and ecological traits influence the direction and strength of butterfly body size shifts? 2) What part do environmental elements like the average temperature before emergence and the land use cover (forests, crops, pastures, and urban areas) play in these dynamics?

We expect that life history traits (phenology, overwintering developmental stage, voltinism, diet breadth, gender) and ecological traits (mobility, thermal tolerance) will have a different effect on butterfly body size. In particular:

1. As warming alters the timing of endotherm hibernation or the spring emergence of holometabolous insects (MacLean et al. 2016), we expect that earlier emergence time will be linked to smaller adult body size because of the shortened development period especially during the warmer years (Fenberg et al. 2016).
2. As some developmental stages are more crucial for determining size at maturity (Kingsolver et al. 2011) we expect that early developmental stages (egg, larva, pupa), more exposed to extreme conditions (cold, dehydration) and directly affected by limitations on energy flow, will show stronger relations to adult size. For example, for a butterfly for which individuals are in the final larval stage during June, it was shown that temperature during this time is a key factor influencing adult size (Fenberg et al. 2016). In addition, for a butterfly for which individuals have a pupal diapause during winter, higher temperatures during the pupal stage drove adults to larger wing sizes (Davies 2019).
3. We expect body size to decrease with the number of generations: univoltine species most often postpone emergence, resulting in a longer larval growth period; hence, size at emergence rises as temperature rises (Horne et al. 2015). In contrast, multivoltine species can enhance voltinism and fit in an extra generation, which causes them to shrink in size since they have to choose between having more generations with less time for growth in each generation and having a longer single generation growth period (Van Dyck et al. 2015).
4. We expect a negative impact of host-plant specificity index to butterfly body size as nutrition specialization can come with evolutionary trade-offs such as investing heavily in adaptations for feeding on a specific plant at the expense of growth and thus body size (Seifert et al. 2022). In addition, we expect mobility traits like larger wings to facilitate efficient movement between fragmented or isolated habitats enabling thus the presence of larger body size individuals.
5. We anticipate that distribution to warmer areas will correspond to high species temperature index (STI) and larger wing length aligning with temperature-size rule and Bergman’s rule (Angilletta and Dunham 2003).

Finally, as sexual size dimorphism is an ongoing debate (Teder 2014, Sõber et al. 2019, Cordeschi et al. 2024) we wanted to investigate if temperature size responses differ between males and females. We used time (years) as a proxy for temperature change over the last century.

## Methods

### Dataset

For our analysis, we utilized metadata exported from two museum collections: (1) the Butterfly Collection of the Swiss Federal Institute of Technology in Zürich (hereafter ETHZ) and (2) the butterfly collection of Goulandris Natural History Museum (hereafter GNHM). Metadata from ETHZ collection included 48,766 specimens representing 639 species collected between 1028 and 2019, while the GNHM collection comprised 6,639 specimens representing 274 species collected between 1917 and 2023.

Excluding specimens with missing information (e.g., collection location or date), a total of 44,316 specimens from ETHZ and 4,582 specimens from GNHM metadata were retained.

We manually measured the length of the right forewing (response variable see below), that is often used as an index for overall body size (Brehm et al. 2019, Minter et al. 2024), on a subset of 17,382 ETHZ specimens and all 5,425 GNHM specimens using ImageJ (https://imagej.nih.gov/ij/index.html).

Additionally, a computer vision tool *(unpublished data*) was applied to measure forewing length for the remaining 44,316 ETHZ specimens. After manual inspection for valid measurements and ventrally pinning of the specimens and further exclusion of duplicates, a total of 23,265 unique specimens remained. To increase the robustness of our sample size we combined the specimens from two methods (manual and automated). The final dataset used for analytical purposes consisted of 31899 unique specimens representing 588 species, with collection dates ranging from 1840 to 2010.

### Life history traits, climatic and spatial data

We compiled 7 species-specific ecological and life-history traits from the literature. We used the European and Maghreb butterfly trait database (Middleton-Welling et al. 2020) for life history traits and CLIMBER database for ecological niche characteristics (Schweiger et al. 2014) (**Table S1**). Our traits selection is narrowed to the ones that are known to affect body size in the light of rising temperatures: (i) hostplant specificity index (HSI) ranges from 0 (highly polyphagous) to 1 (completely monophagous) and provides a quantitative measure of diet, (ii) maximum flight period (FMo_max) gives the maximum number of flight months in a year and drives phenological patterns together with first flight month (FFM), (iii) overwintering stage (OvS: egg, larvae, pupae, adult) is the life history stage in which a species hibernates during winter, (v) mean number of generations per year (Vol_mean), (vi) range size (range.size) as a proxy for mobility that corresponds to the distribution size in Europe and (vii) thermal niche as species temperature index (STI) that averages temperatures of the grid cells where each species is known to occur (Kudrna et al. 2002).

For temperature, we used historical monthly weather data, CHELSAcruts (Karger et al. 2017, Karger and Zimmermann 2018) for the years 1901-1959 at approximately 1 km (30 arc seconds) resolution and WordClim (Fick and Hijmans 2017) for the period 1960 – 2010 at a 2.5 minutes (∼21 km^2^ at the equator) resolution. We combined the two temperature sources and estimated the average value from 1900 to 2010 for two periods: period one (Period1) corresponding to the first three months of a year and period two (Period2) corresponding to the next three months (April-May-June). We used temperatures from months before emergence, when butterflies are in stages before adulthood (egg, early larval, late larval and pupal stages) and the effect of temperature in growth and thus adult fitness is higher (Klockmann and Fischer 2017). We also used Land-Use Harmonization2 (LUH2 v2h) for land-use data (covered period 850-2015) with a spatial resolution of 0.25 × 0.25 degrees (https://luh.umd.edu/data.shtml) and WorldClim 2.1 at a 30 seconds resolution for elevation. Although elevation was not included in the models, in cases where the geometries of raster layers differ, we resampled the CHELSAcruts dataset to match the desired geometry of the WorldClim elevation data, ensuring compatibility for subsequent analyses.

### Modelling

We fitted a generalized additive model to analyze the relationship between forewing length (response variable) and life history traits as well as land use (explanatory variables) using *gam* () function from the mgcv package (Augustin et al. 2012) in R version 4.3.2 (R Core Team 2024). We were aware of the nonlinear relationship between year and response variable from our previous analytical framework and we were not interested in interspecific variation. Therefore, the model was specified as follows:

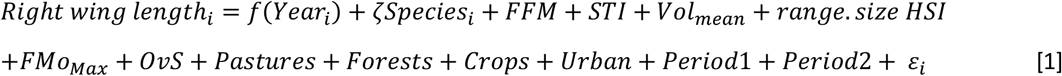

where *f*_1_(*Year*_*i*_) is the smooth term for the continuous predictor*ζSpecies*_*i*_, is the random effect for species to account for random-effect variability in predictions, followed by the parametric terms for life history and ecological traits (*FFM, STI, Vol*_*mean*_, *range*.*size, HSI, FMo*_*Max*_, *Ovs*), land use cover (*Pastures, Forests, Crops, Urban*), estimated mean temperature per year (*Period*1, *Period*2) and *ε*_*i*_ for the Gaussian error term.

We ran a second model including also gender specific trait for a subset of 4,643 individuals, for which sex information was available. We allowed a different smoothing parameter per each gender-specific smoother, suggesting thus a different level of wiggliness (Pedersen et al. 2019). We fit a model with a separate smoother (with its own penalties) for females and males using the following formula:

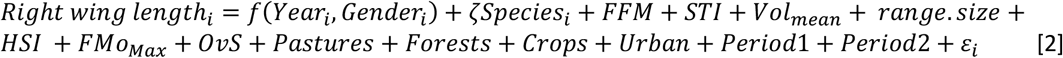

where *f*(*Year*_*i*_, *Gender*_*i*_), is the smooth interaction between the continuous predictor and the grouping factor allowing the effect of Year to vary by gender, while the random effect for species is considered by *ζSpecies*_*i*_. The parametric terms corresponding to life history traits of species (*FFM, STI, Vol*_*mean*_, *range*.*size, HSI, FMo*_*Max*_ + *OvS*) and the ones referring to mean temperature (*Period*1 + *Period*2) as well as the land use cover (*Pastures* + *Forests* + *Crops* + *Urban*) were also included in the model.

Prior to modelling, any missing data for predictor variables at individual level were imputed with chained random forest imputation using the *missRanger* package (Mayer 2024). Model diagnostics, including residual checks and variable importance assessment, were conducted to evaluate model performance and reliability with function *gam*.*check* from mgcv package (Augustin et al. 2012) and function *appraise* from gratia package (Simpson 2024). Plots were made with *parametric_effects* function from gratia package (Simpson 2024).

## Results

The GAM analysis assessed the relationship between butterfly forewing length and various life history, and ecological traits as well as environmental parameters, along with random effects for species (**Figure 1**). The first model explained a substantial portion of the variation in the data (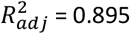, deviance explained = 89.6%). Both phenological traits, i.e., first flight month (FFM) and duration of the flight period (FMo_Max) showed a significant positive effect (β_FFM_ *=* 0.12, *SE* = 0.05, *p* = 0.016; β_FMo_Max_ *=* 0.14, *SE* = 0.03, *p* < 0.001), suggesting that larger body size butterflies appear later in the season and have a longer duration of flight. A positive relationship was found between species temperature index with forewing length (β_STI_ = 0,037, *SE* = 0.01, *p* = 0.006) suggesting that higher STI corresponded to larger sizes.

**Figure 1.**
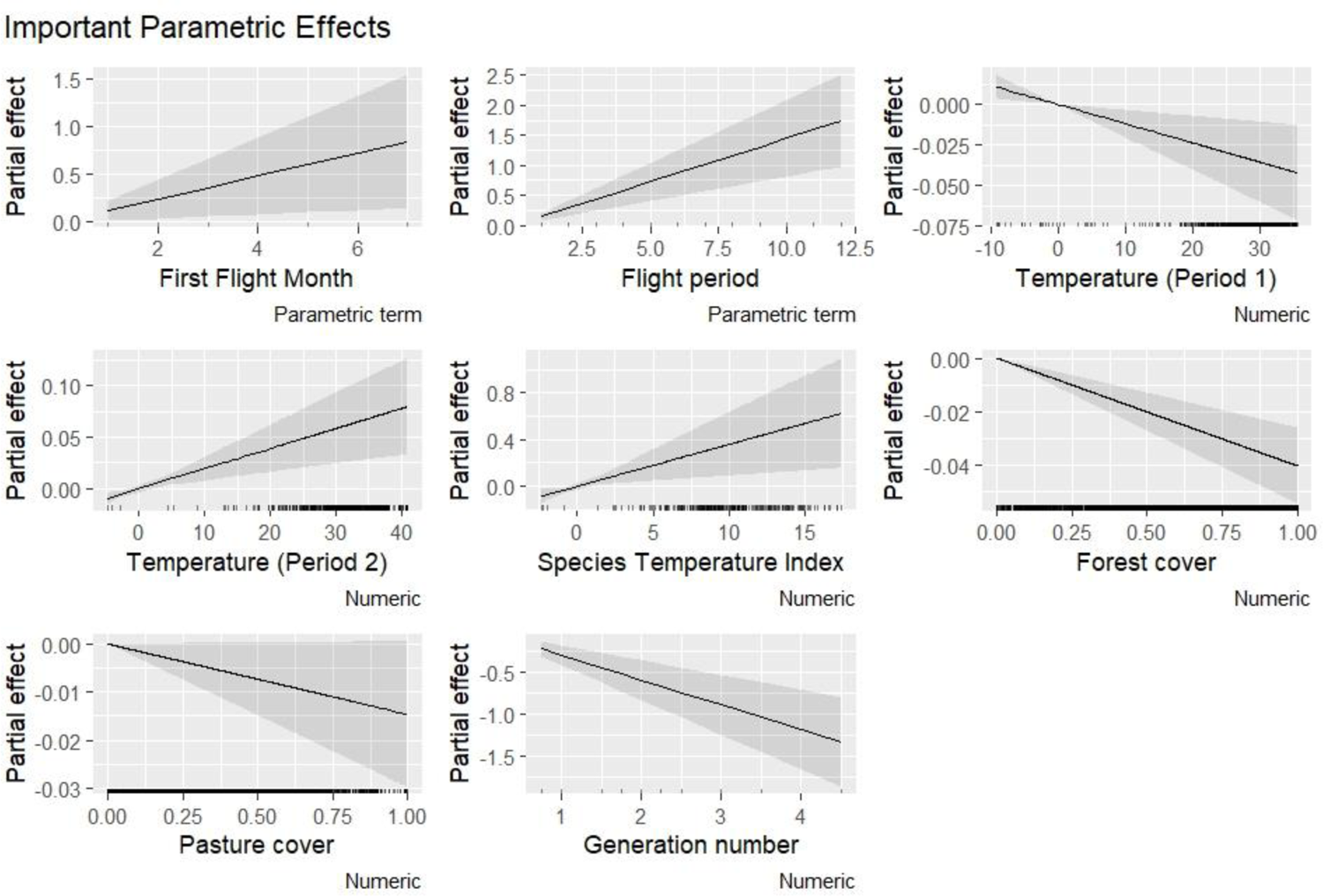
Parametric effects of important predictors on the right forewing length. Each plot represents a different predictor variable from the first GAM model: First Flight Month (FFM); Flight Period (FMo_max); Temperature (Period 1) (Period1); Temperature (Period 2) (Period2); Species Temperature Index (STI); Forest Cover (SumForests); Pasture Cover (SumPastures); Generation Number (Vol_mean). Solid line represents the estimated effect of the predictor on wing length, and the shaded area indicates the 95% confidence interval for the estimated effect, providing a measure of uncertainty around the predictions.

For the number of generations (β_Vol_mean_ = - 0.3, *SE* = 0.06, *p* < 0.001), and the overwintering stage of egg (β_OvSE_ = - 0.88, *SE* = 0.26, *p* = *0*.001) and larvae (β_OvSL_ = - 0.5, *SE* = 0.21, *p* = 0.017) the strong negative effect to wing length indicates a relation to smaller body size. Forest cover (β_Forests_ = - 0.04, *SE* = 0.01, *p < 0*.*001*), pastures (β_Pastures_ = - 0.02, *SE* = 0.01, *p* = 0.036) and mean temperature of the first three months (β_Period1_ = - 0.002, *SE* = 0.0004, *p* < 0.001) also exhibited a negative relationship with wing length. On the contrary, the second period was found to be connected to larger body sizes (β_Period2_ = 0.002, *SE* = 0.001, *p* < 0.001). The environmental effect, however, was very low, close to zero. Model output and diagnostic plots indicated a good fit of the model (**Table S2, Figure S1**).

The second model explained a substantial portion of the variation in the data (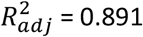, deviance explained = 89.8%) (**Figure 2**). Males’ wing length was found to decrease after 1950, while wide confidence intervals for the first years from 1990 to 1950, suggested that the positive trend might not be valid for this period (**Figure 3**). For females, the linear positive trend (edf = 1) is uncertain for 40 years (1935 – 1975) as confidence intervals included the horizontal line (intercept = 0) (**Figure 3**). All other relationships with parametric terms of the model showed the same patterns we found in our first model with two exceptions: the marginal effect of larval overwintering stage (β_OvSL_ = -0.45, *SE* = 0.28, *p = 0*.*052*) and the lack of any temperature effect (**Table S3**). Model was fully converged after 9 iterations and diagnostic plots indicated that there were no violations of the model assumptions (**Figure S2**).

**Figure 2.**
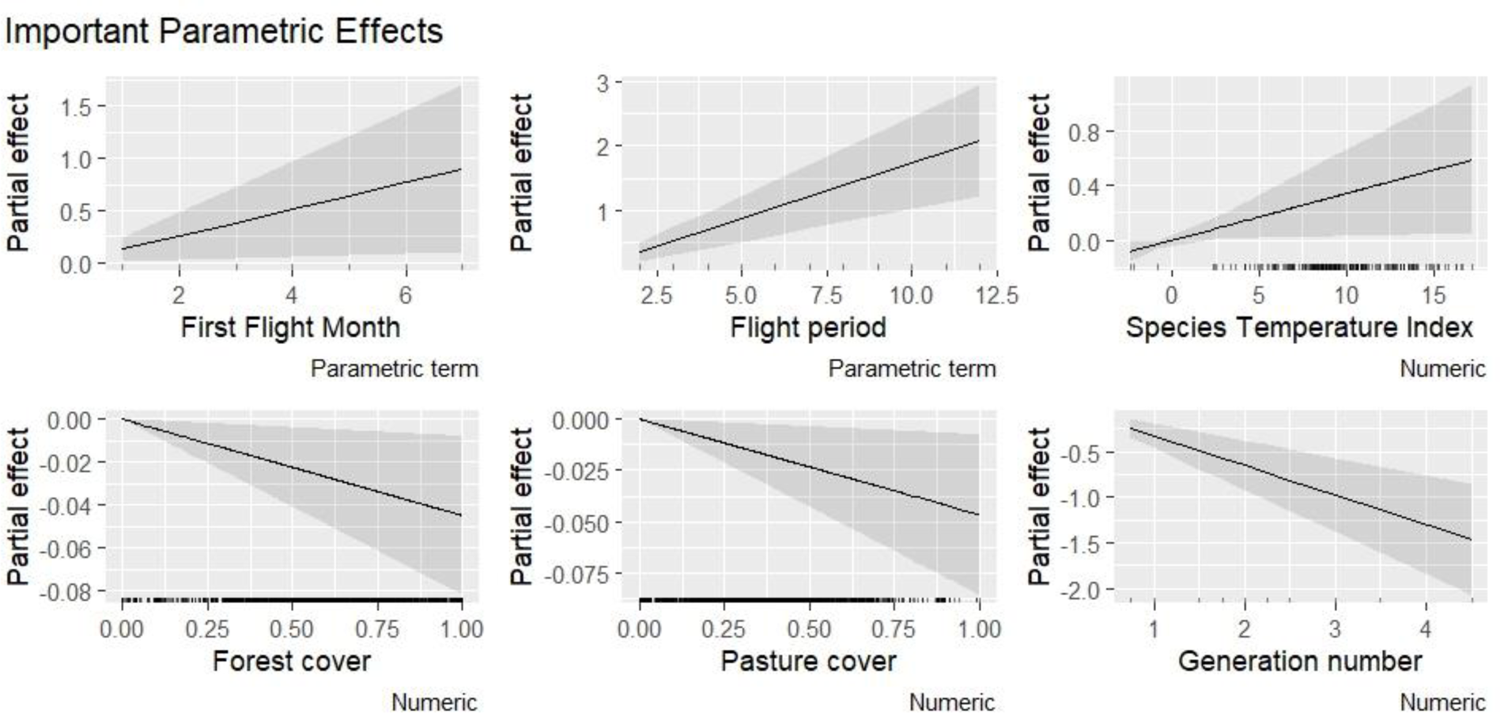
Parametric effects of important predictors on the right forewing length. Each plot represents a different predictor variable from the second GAM model: First Flight Month (FFM); Flight Period (FMo_max); Species Temperature Index (STI); Forest Cover (SumForests); Pasture Cover (SumPastures); Generation Number (Vol_mean). Solid line represents the estimated effect of the predictor on wing length, and the shaded area indicates the 95% confidence interval for the estimated effect, providing a measure of uncertainty around the predictions.

**Figure 3.**
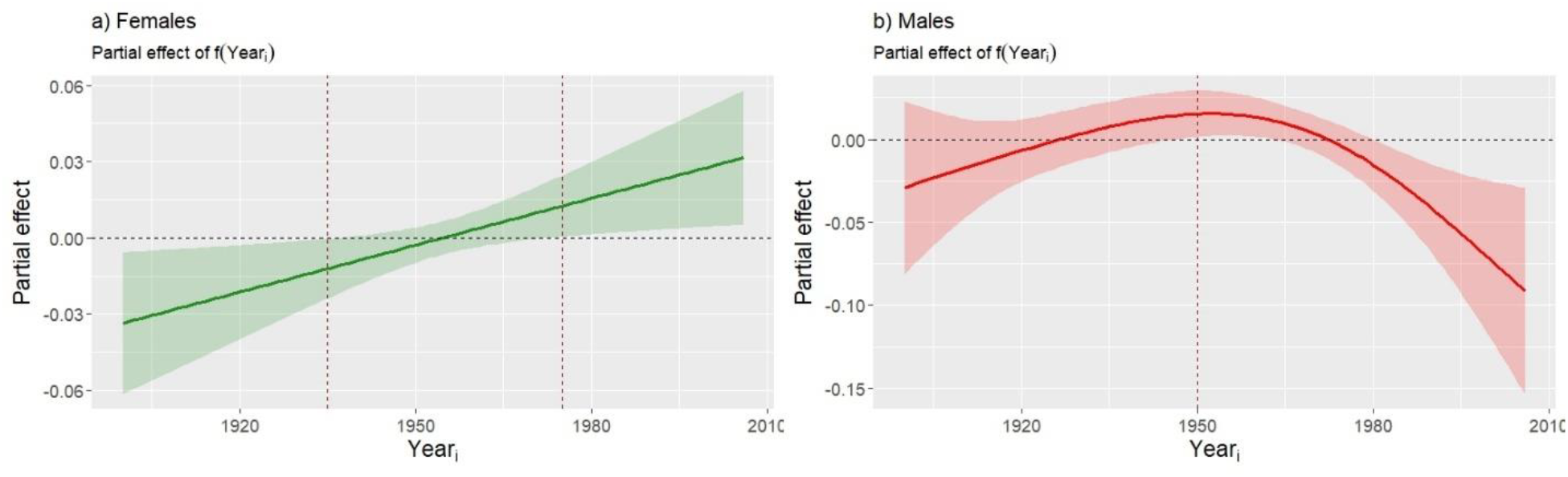
Partial effect plots for the right forewing length and year smooth by gender on the link scale (link = identity). Solid line represents the estimated trend of the smoother year by gender (right: females, left: males) of the predictor on wing length, and the shaded area indicates the 95% confidence intervals for the estimated trend. Red dashed lines indicate important trend, i.e., when horizontal black dashed line is not included inside confidence intervals area.

## Discussion

The adult body size of our butterfly pool was found to differ in life history and ecological-related traits, corresponding largely with our predictions. In addition, this study indicates a change in body size as a response to environmental cues that trigger alternative developmental pathways.

As anticipated, later seasonal appearance was associated with larger body sizes, allowing individuals more time to grow before reaching adulthood. Developmental plasticity in insects, including how larger body sizes require longer growth periods, leads to later emergence (Nylin and Gotthard 1998). For instance, both field data and breeding data of the butterfly *Pararge aegeria* showed that late spring individuals had the longest and largest forewings in both sexes (Van Dyck and Wiklund 2002). Overall, it has been observed that at cooler temperatures, larval growth can be prolonged, sometimes leading to a larger adult size compared to those developing in warmer temperatures (Angilletta et al. 2004).

In Lepidoptera, flight performance is typically linked to morphological changes, including thorax mass, wing size, wing shape, and wing loading (Berwaerts et al. 2002). Overall, larger wings enable butterflies to sustain longer flight periods, likely due to increased energy reserves, though this comes at a cost. For instance, the wing length of diamondback moth is negatively correlated with fecundity, but positively correlated with flight duration ability indicating a trade-off between fecundity and migration (Muhamad et al. 1994). Flight duration capacity, however, depends not only on wing length, but also on energetically costly physiological properties such as muscle mass (Berwaerts et al. 2002) and it is often gender specific, such that males exhibiting longer duration than females (Kingsolver 1983, Gilchrist 1990).

We found a positive relationship between the species temperature index (STI) and forewing length, indicating that species associated with warmer locations exhibited larger body sizes—contrary to our expectations. A possible explanation for the higher STI and larger body sizes could be an adaptive response for better dispersal in fragmented or warmer locations, where mobility is critical for survival. To this extent, (Merckx et al. 2018) found shifts towards larger and thus more mobile individuals when they analyzed macro-moth populations along an urbanization gradient in Belgium. Although the urban heat island effect is expected to drive shifts towards smaller body sizes due to increased metabolic costs, it is also argued that urban settlements are often highly fragmented, which could act as a selecting force for dispersal phenotypes (Merckx et al. 2018).

As expected, we found a negative relationship between the number of generations per year and forewing length suggesting a trade-off between generations and growth. It has been often the case where multivoltine species try to squeeze more generations into the season as a result of elevated temperatures, often with potentially severe fitness consequences, such as developmental traps where an additional generation begins but cannot be completed before winter (Van Dyck et al. 2015, Kerr et al. 2020).

During early stages, butterfly vulnerability to environmental influences is high and can be reflected in shifts to adult body size (Radchuk et al. 2013). Here, we showed that overwintering egg and larva stage (OvS) had a strong negative effect on butterfly body length. This pattern underscores the interplay between developmental stage-specific ecology and the constraints imposed by overwintering strategies (Larsen et al. 2024). For instance, overwintering in earlier developmental stages (egg, larva, pupa) requires surviving on stored energy reserves (Bale and Hayward 2010), unlike adults that might actively forage. Limitations in energy flow could constrain growth, leading to smaller adult sizes. The more recent occurrence of extreme weather temperatures (Meehl and Tebaldi 2004), can also result in reduced body size during overwinter as a trade-off for survival. On the contrary, if cooler microclimatic conditions delay emergence, individuals might have more time to grow before the flight season, reaching a larger final size (Nève and Singer 2008).

The impact of land cover and mean temperatures on butterfly body length reflects a dynamic relationship that shifts over the study period, unlike life history traits that remain rather static over the years. We attribute the observed relationship between forested areas and smaller body size to reduced mobility needed to navigate through this type of habitat and decreased availability of sunlight that forest-dwelling butterflies encounter compared to their counterparts in open or fragmented landscapes (McCoy 2018). As a result, smaller body sizes associated with lower energy and dispersal demands might be favored under that condition. Another parameter that can limit larval growth and thus adult size is reduced access to high quality nutrition as forests typically have lower abundance and diversity of host plants and nectar resources (Stamp 2001). In the light of serious loss of pastures to dynamic urban settlement expansion, we attribute the observed relationship between low pasture cover and larger body size to an adaptive response favoring higher mobility in response to increased habitat fragmentation.

At cooler temperatures, larval growth can be prolonged, sometimes leading to a larger adult size compared to those developing in warmer temperatures (Angilletta et al. 2004). Our findings signify a smaller body size for the first three months of the year (Period 1) but a larger body size for the next three months (Period 2). We argue that rising temperatures over the past century (IPCC 2021) have also affected early spring temperatures, which in turn have led to earlier emergence or hatching of larvae (Davies 2019). As a result, available developmental time for growth before pupation is reduced and individuals may emerge as smaller adults. In addition, warmer early-season temperatures may increase metabolic demands, diverting energy from growth to maintenance. Unlike the constraints of first studied period (January-March), the second period (April-May-June) is likely to offer warmer, more stable conditions that allow larvae to extend their development time accumulating more energy and thus achieving larger adult sizes.

Smaller body sizes in males over the years might be an adaptive response to increasing temperatures, aligning with the size-temperature rule (Verberk et al. 2021). The steep decline in male wing length after 1950 indicates a plastic response to increasing environmental pressures such as elevated temperatures and land use change. The time coincides with increasing CO_2_ concentrations (Keeling 1968, Brewer 2009) and the spread of the Industrial Revolution at mid-20^th^ century. A further temperature increase of up to 4°C by 2100 (IPCC 2021) and further increase of extreme high temperatures (Meehl and Tebaldi 2004), could push butterflies beyond their physiological limits. This risk is particularly pronounced for species living in harsh environments, those more sensitive to environmental changes, and those already near their upper thermal tolerance.

We limit our discussion to positive trend we found for females, during the first (1900-1935) and last years (1980 – 2010), where linear trend seems robust. The positive trend during these periods suggests a more stable selection pressure favoring larger sizes, likely linked to their reproductive role. Larger body size has long been recognized as a key factor enhancing various aspects of organismal performance and evolution (Bonner 2011). Although the “bigger is better” hypothesis holds true in most natural populations, yet reaching and maintaining a larger body size carries its own life history trade-offs such as delayed maturation and increased energy demands. Achieving a larger body size may become increasingly difficult due to anticipated rises in temperature and extreme weather events, as disruptions in trophic interactions can limit available nutritional resources, while earlier emergence times may reduce developmental periods.

### Concluding remarks

Our research shows that life history characteristics, ecological variables, and environmental shifts all affect butterfly body size. We found that larger body sizes are linked to longer flight durations and later seasonal emergence, which is probably an adaptation for the increased mobility required for resource acquisition in fragmented settings. In contrast, smaller sizes in early-season emergence and increasing forest areas reflect trade-offs between growth and energy constraints. Sex-specific patterns suggest that whereas females exhibit more steady trends because of the reproductive benefits of greater sizes, males are more vulnerable to rising temperatures and exhibit a sharp size reduction. Our research emphasizes the critical role of climate change and habitat fragmentation in determining butterfly body size and stresses the need for more research on sexual size dimorphism as anthropogenic climate change might drive butterflies to fail in plastic adjustment of decision-making.

## Acknowledgments

We are grateful to all people for helping us with digitization of the butterfly collection housed in Goulandris Natural History Museum collection, Michael Greeff for facilitating data sharing from the Entomological Collection of Swiss Federal Institute of Technology in Zürich and Christina Kassara for her contribution in spatial analysis.

## Funding

The research project was supported by the Hellenic Foundation for Research and Innovation (H.F.R.I.) under the “3rd Call for H.F.R.I. Research Projects to support Post-Doctoral Researchers” (Project Number 7191).

## Author contributions

K.Z. conceived and performed the study. C.M.DM., M.R.L.W., facilitated the data use from ETHZ collection. K.Z. secured funding and V.K. supervised the project. Data analysis conducted by K.Z. The manuscript was originally drafted by K.Z. and E.K., B.J.S., O.S., C.M.DM, M.R.L.W. and V.K. contributed to the interpretation of results, manuscript revisions, and approved the final version for submission.

## Code availability

The R-script of the analysis is available at Zenodo (DOI: 10.5281/zenodo.14803566)

## Notes

https://doi.org/10.5281/zenodo.14803566

### Competing Interest Statement

The authors have declared no competing interest.

## References

Altermatt, F. 2010. Tell me what you eat and I’ll tell you when you fly: diet can predict phenological changes in response to climate change. Ecology Letters 13:1475–1484.

Altermatt, F. 2012. Temperature-related shifts in butterfly phenology depend on the habitat. Global Change Biology 18:2429–2438.

Angilletta, J. Michael J., and Arthur E. Dunham. 2003. The Temperature-Size Rule in Ectotherms: Simple Evolutionary Explanations May Not Be General. The American Naturalist 162:332–342.

Angilletta, M. J., Jr., T. D. Steury, and M. W. Sears. 2004. Temperature, Growth Rate, and Body Size in Ectotherms: Fitting Pieces of a Life-History Puzzle1. Integrative and Comparative Biology 44:498–509.

Augustin, N. H., E.-A. Sauleau, and S. N. Wood. 2012. On quantile quantile plots for generalized linear models. Computational Statistics & Data Analysis 56:2404–2409.

Bale, J. S., and S. A. L. Hayward. 2010. Insect overwintering in a changing climate. Journal of Experimental Biology 213:980–994.

Berwaerts, K., H. Van Dyck, and P. Aerts. 2002. Does flight morphology relate to flight performance? An experimental test with the butterfly Pararge aegeria. Functional Ecology 16:484–491.

Bonner, J. T. 2011. Why size matters: From bacteria to blue whales. Why Size Matters: From Bacteria to Blue Whales.

Brehm, G., D. Zeuss, and R. K. Colwell. 2019. Moth body size increases with elevation along a complete tropical elevational gradient for two hyperdiverse clades. Ecography 42:632–642.

Brewer, P. G. 2009. The influence of david keeling on oceanic CO2 measurements. Geophysical Monograph Series 183:37–48.

Cordeschi, G., D. Canestrelli, and D. Porretta. 2024. Sex-biased phenotypic plasticity affects sexual dimorphism patterns under changing environmental conditions. Scientific Reports 14:892.

Davies, W. J. 2019. Multiple temperature effects on phenology and body size in wild butterflies predict a complex response to climate change. Ecology 100:e02612.

Fenberg, P. B., A. Self, J. R. Stewart, R. J. Wilson, and S. J. Brooks. 2016. Exploring the universal ecological responses to climate change in a univoltine butterfly. J Anim Ecol 85:739–748.

Fick, S. E., and R. J. Hijmans. 2017. WorldClim 2: new 1-km spatial resolution climate surfaces for global land areas. International Journal of Climatology 37:4302–4315.

Fischer, K., and K. Fiedler. 2002. Life-history plasticity in the butterfly Lycaena hippothoe: local adaptations and trade-offs. Biological Journal of the Linnean Society 75:173–185.

Garcia, R. A., M. Cabeza, C. Rahbek, and M. B. Araújo. 2014. Multiple Dimensions of Climate Change and Their Implications for Biodiversity. Science 344:1247579.

Gilchrist, G. 1990. The consequences of sexual dimorphism in body size for butterfly flight and thermoregulation. Functional Ecology:475–487.

Hill, G. M., A. Y. Kawahara, J. C. Daniels, C. C. Bateman, and B. R. Scheffers. 2021. Climate change effects on animal ecology: butterflies and moths as a case study. Biological Reviews 96:2113–2126.

Hoffmann, A. A., and C. M. Sgrò. 2011. Climate change and evolutionary adaptation. Nature 470:479–485.

Horne, C. R., A. G. Hirst, and D. Atkinson. 2015. Temperature-size responses match latitudinal-size clines in arthropods, revealing critical differences between aquatic and terrestrial species. Ecology Letters 18:327–335.

IPCC. 2021. Climate Change 2021: The Physical Science Basis. “In Contribution of Working Group I to the Sixth Assessment Report of the Intergovernmental Panel on Climate Change, edited by V. Masson-Delmotte, P. Zhai, A. Pirani, S. L. Connors, C. Péan, S. Berger, N. Caud, Y. Chen, L. Goldfarb, M. I. Gomis, M. Huang, K. Leitzell, E. Lonnoy, J. B. R. Matthews, T. K. Maycock, T. Waterfield, O. Yelekçi, R. Yu, and B. Zhou. Cambridge and New York, NY: Cambridge University Press.

Karger, D. N., O. Conrad, J. Böhner, T. Kawohl, H. Kreft, R. W. Soria-Auza, N. E. Zimmermann, H. P. Linder, and M. Kessler. 2017. Climatologies at high resolution for the earth’s land surface areas. Scientific Data 4:170122.

Karger, D. N., and N. E. Zimmermann. 2018. CHELSAcruts - High resolution temperature and precipitation timeseries for the 20th century and beyond. EnviDat.

Karl, I., R. Stoks, M. De Block, S. A. Janowitz, and K. Fischer. 2011. Temperature extremes and butterfly fitness: conflicting evidence from life history and immune function. Global Change Biology 17:676–687.

Keeling, C. D. 1968. Carbon dioxide in surface ocean waters: 4. Global distribution. Journal of Geophysical Research (1896-1977) 73:4543–4553.

Kerr, N. Z., T. Wepprich, F. S. Grevstad, E. B. Dopman, F. S. Chew, and E. E. Crone. 2020. Developmental trap or demographic bonanza? Opposing consequences of earlier phenology in a changing climate for a multivoltine butterfly. Global Change Biology 26:2014–2027.

Kingsolver, J. G. 1983. Thermoregulation and flight in Colias butterflies: elevational patterns and mechanistic limitations. Ecology 64:534–545.

Kingsolver, J. G., H. Arthur Woods, L. B. Buckley, K. A. Potter, H. J. MacLean, and J. K. Higgins. 2011. Complex Life Cycles and the Responses of Insects to Climate Change. Integrative and Comparative Biology 51:719–732.

Kingsolver, J. G., S. E. Diamond, and L. B. Buckley. 2013. Heat stress and the fitness consequences of climate change for terrestrial ectotherms. Functional Ecology 27:1415–1423.

Klockmann, M., and K. Fischer. 2017. Effects of temperature and drought on early life stages in three species of butterflies: Mortality of early life stages as a key determinant of vulnerability to climate change? Ecology and Evolution 7:10871–10879.

Kudrna, O., N. Deutschland, G. f. Schmetterlingsschutz, and M. E. Butterflies. 2002. The Distribution Atlas of European Butterflies. Apollo Books.

Larsen, E. A., M. W. Belitz, G. J. Di Cecco, J. Glassberg, A. H. Hurlbert, L. Ries, and R. P. Guralnick. 2024. Overwintering strategy regulates phenological sensitivity and consequences for ecological services in a clade of temperate North American insects. Functional Ecology 38:1075–1088.

MacLean, H. J., J. G. Kingsolver, and L. B. Buckley. 2016. Historical changes in thermoregulatory traits of alpine butterflies reveal complex ecological and evolutionary responses to recent climate change. Climate Change Responses 3:13.

Mayer, M. 2024. missRanger: Fast Imputation of Missing Values. R package version 2.6.1, https://mayer79.github.io/missRanger/, https://github.com/mayer79/missRanger.

McCoy, S. 2018. Butterfly wing shape variation among habitats and their phylogenetic relationships, June 2018”. Monteverde Institute: Tropical Ecology and Conservation. 204. https://digitalcommons.usf.edu/tropical_ecology/204.

Meehl, G. A., and C. Tebaldi. 2004. More intense, more frequent, and longer lasting heat waves in the 21st century. Science 305:994–997.

Merckx, T., A. Kaiser, and H. Van Dyck. 2018. Increased body size along urbanization gradients at both community and intraspecific level in macro-moths. Global Change Biology 24:3837–3848.

Middleton-Welling, J., L. Dapporto, E. García-Barros, M. Wiemers, P. Nowicki, E. Plazio, S. Bonelli, M. Zaccagno, M. Šašić, J. Liparova, O. Schweiger, A. Harpke, M. Musche, J. Settele, R. Schmucki, and T. Shreeve. 2020. A new comprehensive trait database of European and Maghreb butterflies, Papilionoidea. Scientific Data 7:351.

Minter, M., K. K. Dasamahapatra, M. D. Morecroft, C. D. Thomas, and J. K. Hill. 2024. Reduced size in a montane butterfly at its warm range boundaries. Ecological Entomology 49:983–988.

Muhamad, O., R. Tsukuda, Y. Oki, K. Fujisaki, and F. Nakasuji. 1994. Influences of wild crucifers on life history traits and flight ability of the diamondback moth,Plutella xylostella (Lepidoptera: Yponomeutidae). Researches on Population Ecology 36:53–62.

Nève, G., and M. C. Singer. 2008. Protandry and postandry in two related butterflies: conflicting evidence about sex-specific trade-offs between adult size and emergence time. Evolutionary Ecology 22:701–709.

Nylin, S., and K. Gotthard. 1998. Plasticity in life-history traits. Annu Rev Entomol 43:63–83.

Oliver, T. H., and M. D. Morecroft. 2014. Interactions between climate change and land use change on biodiversity: attribution problems, risks, and opportunities. WIREs Climate Change 5:317–335.

Parmesan, C. 2003. CHAPTER 24. Butterflies as Bioindicators for Climate Change Effects. Pages 541–560 in L. B. Carol, B. W. Ward, and R. E. Paul, editors. Butterflies. University of Chicago Press, Chicago.

Pedersen, E. J., D. L. Miller, G. L. Simpson, and N. Ross. 2019. Hierarchical generalized additive models in ecology: an introduction with mgcv. PeerJ 7:e6876.

R Core Team, R. 2024. “R: A language and environment for statistical computing. R Foundation for Statistical Computing, Vienna, Austria. URL https://www.R-project.org/.“.

Radchuk, V., C. Turlure, and N. Schtickzelle. 2013. Each life stage matters: the importance of assessing the response to climate change over the complete life cycle in butterflies. Journal of Animal Ecology 82:275–285.

Scheffers, B. R., L. De Meester, T. C. L. Bridge, A. A. Hoffmann, J. M. Pandolfi, R. T. Corlett, S. H. M. Butchart, P. Pearce-Kelly, K. M. Kovacs, D. Dudgeon, M. Pacifici, C. Rondinini, W. B. Foden, T. G. Martin, C. Mora, D. Bickford, and J. E. M. Watson. 2016. The broad footprint of climate change from genes to biomes to people. Science 354:aaf7671.

Schweiger, O., A. Harpke, M. Wiemers, and J. Settele. 2014. CLIMBER: Climatic niche characteristics of the butterflies in Europe. ZooKeys 367:65–84.

Seifert, C. L., P. Strutzenberger, and K. Fiedler. 2022. Ecological specialisation and range size determine intraspecific body size variation in a speciose clade of insect herbivores. Oikos 2022:e09338.

Simpson, G. 2024. gratia: Graceful ggplot-Based Graphics and Other Functions for GAMs Fitted using mgcv. R package version 0.9.2, https://gavinsimpson.github.io/gratia/.

Singer, M. C., and C. Parmesan. 2010. Phenological asynchrony between herbivorous insects and their hosts: signal of climate change or pre-existing adaptive strategy? Philosophical Transactions of the Royal Society B: Biological Sciences 365:3161–3176.

Sõber, V., S. L. Sandre, T. Esperk, T. Teder, and T. Tammaru. 2019. Ontogeny of sexual size dimorphism revisited: Females grow for a longer time and also faster. PLoS One 14:e0215317.

Stamp, N. E. 2001. Effects of prey quantity and quality on predatory wasps. Ecological Entomology 26:292–301.

Teder, T. 2014. Sexual size dimorphism requires a corresponding sex difference in development time: a meta-analysis in insects. Functional Ecology 28:479–486.

Van Dyck, H., D. Bonte, R. Puls, K. Gotthard, and D. Maes. 2015. The lost generation hypothesis: could climate change drive ectotherms into a developmental trap? Oikos 124:54–61.

Van Dyck, H., and C. Wiklund. 2002. Seasonal butterfly design: morphological plasticity among three developmental pathways relative to sex, flight and thermoregulation. Journal of Evolutionary Biology 15:216–225.

Verberk, W. C. E. P., D. Atkinson, K. N. Hoefnagel, A. G. Hirst, C. R. Horne, and H. Siepel. 2021. Shrinking body sizes in response to warming: explanations for the temperature–size rule with special emphasis on the role of oxygen. Biological Reviews 96:247–268.

Zografou, K., M. T. Swartz, G. C. Adamidis, V. P. Tilden, E. N. McKinney, and B. J. Sewall. 2021. Species traits affect phenological responses to climate change in a butterfly community. Scientific Reports 11:3283.

