## Supporting Information for "Butterfly body size shrinkage: the impact of ecological traits across varied environments"

This file includes supplementary tables, figures, and additional analyses that support the conclusions of the main manuscript. It is intended to provide transparency and reproducibility for the reported findings.

**Contents:**

Table S1-S3

Figure S1-S3

For any questions regarding this material, please contact []

**Table S1.** List with life history traits, ecological traits, land use cover data and temperature data we included in our modelling approach.

|  | **Explanatory variables** | **Description** | **Range/Levels** | **Source** |
| --- | --- | --- | --- | --- |
| Life history traits | HSI | Hostplant Specificity Index | 0 - 1 | Middleton-Welling et al. 2020 |
|  | FMo_max | Maximum flight period | 1 - 12 | Middleton-Welling et al. 2021 |
|  | FFM | First Flight Month | 1 - 7 | Middleton-Welling et al. 2022 |
|  | OvS | Overwintering stage | egg; larvae; pupae; adult | Middleton-Welling et al. 2023 |
|  | Vol_mean | Mean annual generations | 0.75 - 4.5 | Middleton-Welling et al. 2024 |
| Ecological traits | range.size | Geographic distribution size | 1 - 1476 grid cells | Schweiger et al. 2014 |
|  | STI | Species Temperature Index | from -2 to 18 in Celcius | Schweiger et al. 2015 |
| Climatic & Spatial data | Tmax_chelsa | monthly average maximum temperature | from 1901 to 1959 in Celcius | <https://chelsa-climate.org/chelsacruts/> |
|  | Tmax_wc | Monthly average maximum temperature | from 1960 to 2010 in Celcius | <https://www.worldclim.org/data/monthlywth.html> |
|  | urban | urban land | 0 - 1 | [Land-Use Harmonization2 https://luh.umd.edu/data.shtml](https://luh.umd.edu/data.shtml) |
|  | Pastures | Sum of managed pastures and rangelands | 0 - 1 | [Land-Use Harmonization2 https://luh.umd.edu/data.shtml](https://luh.umd.edu/data.shtml) |
|  | Crops | Sum of annual crops, perennial crops, annual crops, perennial crops and nitrogen-fixing crops | 0 - 1 | [Land-Use Harmonization2 https://luh.umd.edu/data.shtml](https://luh.umd.edu/data.shtml) |
|  | Forests | Sum of forested primary land, non-forested primary land, potentially forested secondary land and potentially non-forested secondary land | 0 - 1 | [Land-Use Harmonization2 https://luh.umd.edu/data.shtml](https://luh.umd.edu/data.shtml) |
|  | Elevation | Elevation in meters | 30 seconds resolution | WorldClim 2.1 |

**Table S2.** Output of our first model where forewing length is a function of two smoothers: a TPRS (thin plate regression spline) single common global smoother of year and a random effect of species. The parametric part of the function includes butterfly life history and ecological traits (FFM, STI, Vol_mean, range.size, HSI, Fmo_Max, OvS), land use cover (Pastures, Forests, Crops, Urban areas) and mean temperature for January-February-March (Period1) as well as temperature for April-May-June (Period2). The response variable (Wing length) is the length of the forewing defines as the span from the point where it connects to the thorax to the wing tip.

| Response variable | Parametric coefficients | Estimate | Std.Error | t value | Pr(>\|t\|) |
| --- | --- | --- | --- | --- | --- |
| Wing length | (Intercept) | 1.622 | 0.426 | 3.811 | **<0.001** |
|  | FFM | 0.123 | 0.051 | 2.417 | **0.016** |
|  | STI | 0.037 | 0.014 | 2.750 | **0.006** |
|  | Vol_mean | -0.297 | 0.061 | -4.898 | **<0.001** |
|  | range.size | 0.000 | 0.000 | -0.705 | 0.481 |
|  | HSI | -0.012 | 0.095 | -0.131 | 0.896 |
|  | FMo_Max | 0.144 | 0.033 | 4.399 | **<0.001** |
|  | OvSE | -0.878 | 0.256 | -3.427 | **0.001** |
|  | OvSEL | -0.256 | 0.259 | -0.989 | 0.323 |
|  | OvSELP | -0.758 | 0.672 | -1.127 | 0.260 |
|  | OvSELPA | -0.249 | 0.277 | -0.897 | 0.370 |
|  | OvSL | -0.501 | 0.210 | -2.380 | **0.017** |
|  | OvSLP | -0.504 | 0.266 | -1.891 | 0.059 |
|  | OvSP | -0.406 | 0.216 | -1.880 | 0.060 |
|  | SumPastures | -0.016 | 0.008 | -2.091 | **0.036** |
|  | SumForests | -0.040 | 0.007 | -5.521 | **<0.001** |
|  | SumCrops | -0.067 | 0.045 | -1.474 | 0.140 |
|  | urban | 0.173 | 0.362 | 0.478 | 0.633 |
|  | Per1_chel | -0.002 | 0.000 | -3.993 | **<0.001** |
|  | Per2_chel | 0.002 | 0.001 | 4.125 | **<0.001** |
|  | Smooth terms | edf | Ref.df | F | p-value |
|  | s(Year) | 3.491 | 3.856 | 24.150 | **<0.001** |
|  | s(Species) | 305.693 | 313.000 | 571.690 | **<0.001** |
| Explained deviance = 89.6% | |  |  |  |  |

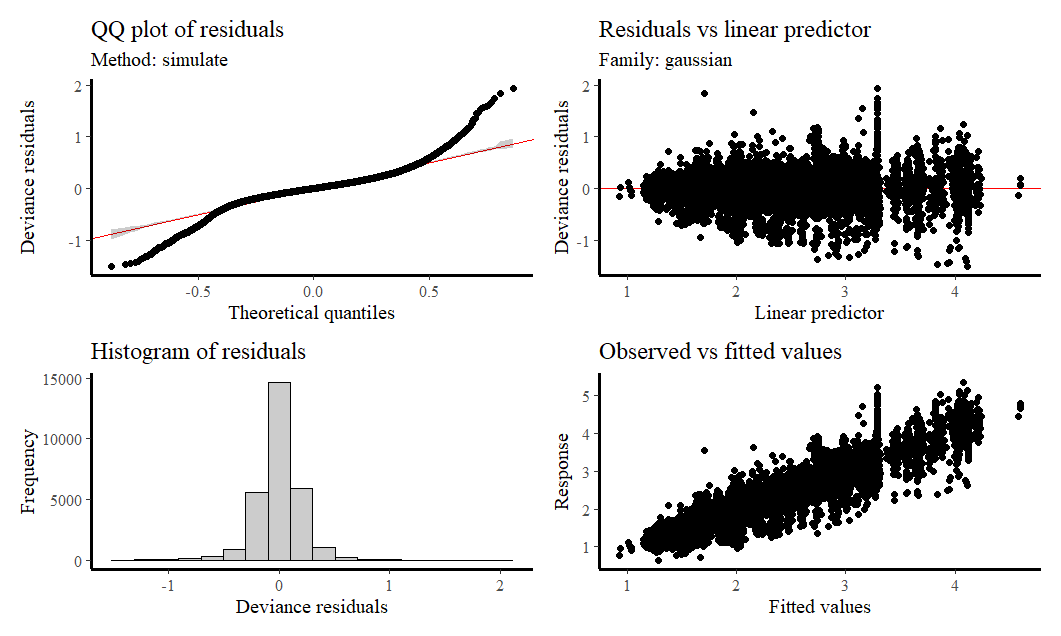

**Figure S1**. The diagnostic plots generated from the first GAM indicated that there were no violations of the model assumptions (n = 28999). The model was fully converged after 6 iterations.

**Table S3.** Output of our second model where forewing length is a function of two smoothers: a separate year smoother per gender and a random effect of species. The parametric part of the function includes butterfly life history and ecological traits (FFM, STI, Vol_mean, range.size, HSI, Fmo_Max, OvS), land use cover (Pastures, Forests, Crops, Urban areas) and mean temperature for January-February-March (Period1) as well as temperature for April-May-June (Period2). The response variable (Wing length) is the length of the forewing defines as the span from the point where it connects to the thorax to the wing tip.

| Response variable | Parametric coefficients | Estimate | Std.Error | t value | Pr(>\|t\|) |
| --- | --- | --- | --- | --- | --- |
| Wing length | (Intercept) | 1.47 | 0.49 | 3.03 | **0.002** |
|  | FFM | 0.13 | 0.06 | 2.19 | **0.028** |
|  | STI | 0.03 | 0.02 | 2.08 | **0.038** |
|  | Vol_mean | -0.33 | 0.07 | -4.72 | **<0.001** |
|  | range.size | 0.00 | 0.00 | -0.98 | 0.329 |
|  | HSI | 0.07 | 0.14 | 0.46 | 0.646 |
|  | FMo_Max | 0.17 | 0.04 | 4.76 | **<0.001** |
|  | OvSE | -0.77 | 0.28 | -2.73 | **0.006** |
|  | OvSEL | -0.23 | 0.28 | -0.81 | 0.418 |
|  | OvSELP | -0.68 | 0.69 | -0.99 | 0.322 |
|  | OvSELPA | -0.14 | 0.31 | -0.46 | 0.647 |
|  | OvSL | -0.45 | 0.23 | -1.94 | 0.052 |
|  | OvSLP | -0.45 | 0.28 | -1.60 | 0.110 |
|  | OvSP | -0.37 | 0.23 | -1.58 | 0.114 |
|  | urban | 0.52 | 0.86 | 0.61 | 0.544 |
|  | SumForests | -0.05 | 0.02 | -2.48 | **0.013** |
|  | SumPastures | -0.05 | 0.02 | -2.53 | **0.012** |
|  | SumCrops | -0.11 | 0.11 | -0.99 | 0.322 |
|  | Per1_chel | 0.00 | 0.00 | -0.63 | 0.527 |
|  | Per2_chel | 0.00 | 0.00 | 1.77 | 0.077 |
|  | Smooth terms | edf | Ref.df | F | p-value |
|  | s(Year):SexFemale | 1 | 1 | 5.77 | **0.016** |
|  | s(Year):SexMale | 2.70 | 3.24 | 3.89 | **0.008** |
|  | s(Species) | 249.542 | 259 | 94.795 | **<0.001** |
| Explained deviance = 89.8% | |  |  |  |  |

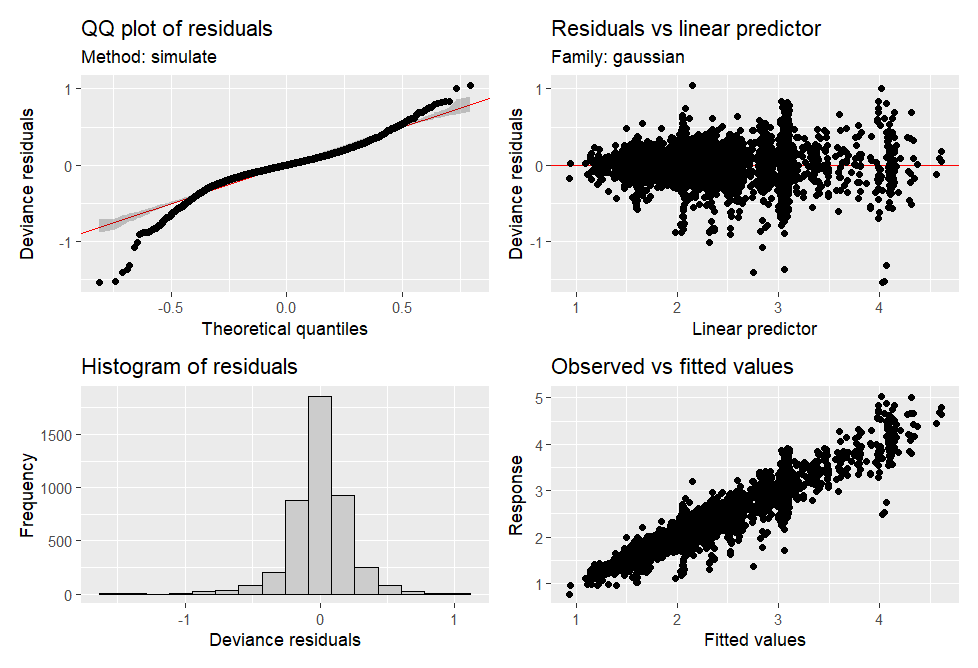

**Figure S2**. The diagnostic plots generated from the first GAM indicated that there were no violations of the model assumptions (n = 4357). Model was fully converged after 9 iterations.

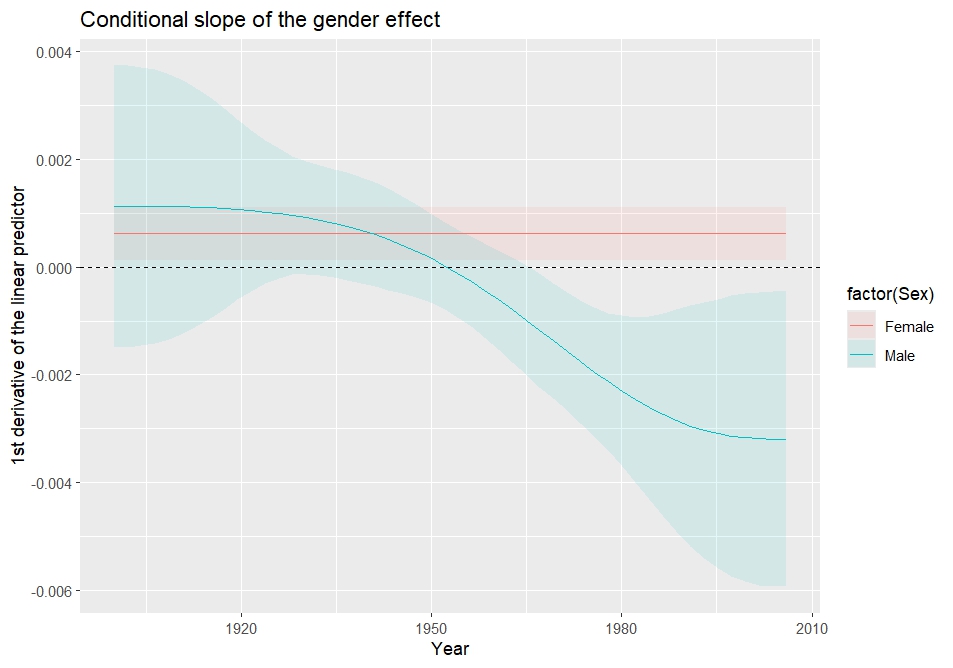

**Figure S3. Slopes of the fitted function over time as calculated by the first derivatives of smoother year per Gender using *plot_slopes ()* function. Both females and males show a significant trend. Males’ slope after 1950 remains below zero justifying the decreasing trend we showed for that period, while females’ slope is entirely above zero line justifying the positive trend we found.**
